# Extracellular cysteine disulfide bond break at Cys122 disrupts PIP_2_-dependent Kir2.1 channel function and leads to arrhythmias in Andersen-Tawil Syndrome

**DOI:** 10.1101/2023.06.07.544151

**Authors:** Francisco M. Cruz, Álvaro Macías, Ana I. Moreno-Manuel, Lilian K. Gutiérrez, María Linarejos Vera-Pedrosa, Isabel Martínez-Carrascoso, Patricia Sánchez Pérez, Juan Manuel Ruiz Robles, Francisco J Bermúdez-Jiménez, Aitor Díaz-Agustín, Fernando Martínez de Benito, Salvador Arias Santiago, Aitana Braza-Boils, Mercedes Martín-Martínez, Marta Gutierrez-Rodríguez, Juan A. Bernal, Esther Zorio, Juan Jiménez-Jaimez, José Jalife

## Abstract

**Background:** Andersen-Tawil Syndrome Type 1 (ATS1) is a rare heritable disease caused by mutations in the strong inwardly rectifying K^+^ channel Kir2.1. The extracellular Cys122-to-Cys154 disulfide bond in the Kir2.1 channel structure is crucial for proper folding, but has not been associated with correct channel function at the membrane. We tested whether a human mutation at the Cys122-to-Cys154 disulfide bridge leads to Kir2.1 channel dysfunction and arrhythmias by reorganizing the overall Kir2.1 channel structure and destabilizing the open state of the channel.

**Methods and Results:** We identified a Kir2.1 loss-of-function mutation in Cys122 (c.366 A>T; p.Cys122Tyr) in a family with ATS1. To study the consequences of this mutation on Kir2.1 function we generated a cardiac specific mouse model expressing the Kir2.1^C122Y^ mutation. Kir2.1^C122Y^ animals recapitulated the abnormal ECG features of ATS1, like QT prolongation, conduction defects, and increased arrhythmia susceptibility. Kir2.1^C122Y^ mouse cardiomyocytes showed significantly reduced inward rectifier K^+^ (I_K1_) and inward Na^+^ (I_Na_) current densities independently of normal trafficking ability and localization at the sarcolemma and the sarcoplasmic reticulum. Kir2.1^C122Y^ formed heterotetramers with wildtype (WT) subunits. However, molecular dynamic modeling predicted that the Cys122-to-Cys154 disulfide-bond break induced by the C122Y mutation provoked a conformational change over the 2000 ns simulation, characterized by larger loss of the hydrogen bonds between Kir2.1 and phosphatidylinositol-4,5-bisphosphate (PIP_2_) than WT. Therefore, consistent with the inability of Kir2.1^C122Y^ channels to bind directly to PIP_2_ in bioluminescence resonance energy transfer experiments, the PIP_2_ binding pocket was destabilized, resulting in a lower conductance state compared with WT. Accordingly, on inside-out patch-clamping the C122Y mutation significantly blunted Kir2.1 sensitivity to increasing PIP_2_ concentrations.

**Conclusion:** The extracellular Cys122-to-Cys154 disulfide bond in the tridimensional Kir2.1 channel structure is essential to channel function. We demonstrated that ATS1 mutations that break disulfide bonds in the extracellular domain disrupt PIP_2_-dependent regulation, leading to channel dysfunction and life-threatening arrhythmias.

**CLINICAL PERSPECTIVE:** *NOVELTY AND SIGNIFICANCE:* **What is known?** - Andersen-Tawil Syndrome Type 1 (ATS1) is a rare arrhythmogenic disease caused by loss-of-function mutations in *KCNJ2*, the gene encoding the strong inward rectifier potassium channel Kir2.1 responsible for I_K1_.
- Extracellular Cys_122_ and Cys_154_ form an intramolecular disulfide bond that is essential for proper Kir2.1 channel folding but not considered vital for channel function.
- Replacement of Cys_122_ or Cys_154_ residues in the Kir2.1 channel with either alanine or serine abolished ionic current in *Xenopus laevis* oocytes. **What new information does this article contribute?** - We generated a mouse model that recapitulates the main cardiac electrical abnormalities of ATS1 patients carrying the C122Y mutation, including prolonged QT interval and life-threatening ventricular arrhythmias.
- We demonstrate for the first time that a single residue mutation causing a break in the extracellular Cys122-to-Cys154 disulfide-bond leads to Kir2.1 channel dysfunction and arrhythmias in part by reorganizing the overall Kir2.1 channel structure, disrupting PIP2-dependent Kir2.1 channel function and destabilizing the open state of the channel.
- Defects in Kir2.1 energetic stability alter the functional expression of the voltage-gated cardiac sodium channel Nav1.5, one of the main Kir2.1 interactors in the macromolecular channelosome complex, contributing to the arrhythmias.
- The data support the idea that susceptibility to arrhythmias and SCD in ATS1 are specific to the type and location of the mutation, so that clinical management should be different for each patient.
- Altogether, the results may lead to the identification of new molecular targets in the future design of drugs to treat a human disease that currently has no defined therapy.

## Introduction

Andersen-Tawil syndrome type 1 (ATS1) is a rare, inheritable autosomal dominant disease caused by loss-of-function mutations in the *KCNJ2* gene, which codes the strong inward rectifier potassium channel Kir2.1.^1,2^ Kir2.1 is ubiquitously expressed throughout the human body and ATS1 mutations predispose patients to a triad of alterations including periodic paralysis, dysmorphias, and arrhythmias that can lead to sudden cardiac death (SCD)^3,4^ by mechanisms that remain unclear.^5^ In the heart, Kir2.1 is responsible for the inward rectifier K^+^ current (I_K1_),^6^ which plays a central role in the maintenance of the resting membrane potential (RMP) and the final phase of action potential (AP) repolarization.^7^ Therefore, loss-of-function mutations in Kir2.1 lead to a substantial decrease in I_K1_, with consequent membrane depolarization at rest, as well as AP duration (APD) and QT interval prolongation.^8^ Normal Kir2.1 channel function requires agonist phosphatidylinositol-4, 5-bisphospate (PIP_2_) interactions, which stabilizes the G-loop in the open state. Defects in PIP_2_ binding are a major pathophysiologic mechanism underlying the loss-of-function phenotype for several ATS1 associated mutations.^5,9–11^

The primary structure of the human Kir2.1 channel comprises a total of thirteen cysteine (Cys) residues distributed along each monomer. Cys residues are uniquely reactive providing the ability to form disulfide bonds.^12^ They contribute to the structural stability of proteins while being key target sites for redox related processes.^13^ Thus, Cys mutations may affect the tridimensional structure of the channel and alter its function. Seven Cys are expected to be distributed in the Kir2.1 channel N- and C-terminus regions, but mutation in most of them have not been shown to significantly affect the single-channel conductance nor the channel open probability.^14^ However, mutating Cys_76_ and Cys_311_ to polar or charged residues modulated the interaction between Kir2.1 and PIP_2_, and resulted in either an absence of channel activity or a decrease in open probability.^14^ Similarly, class Ic antiarrhythmic drugs have been shown to bind to the Cys_311_ residue of the Kir2.1 channel, and to reduce the polyamine-induced inward rectification increasing the outward I_K1_.^15,16^ Four Cys residues are located in the channel transmembrane segment TM1 (Cys_89_ and Cys_101_), the pore (Cys_149_) and TM2 regions (Cys_169_). Importantly, the remaining two Cys, Cys_122_ and Cys_154_, are located at extracellular space positions absolutely conserved across the inward rectifier family,^17^ and form a disulfide bond crucial for channel assembly.^12,18,19^ However, the Cys122-to-Cys154 disulfide bridge has not been considered essential for normal Kir2.1 function once the channel has been formed.^12^

Here we report on an ATS1 family with a novel Kir2.1 loss-of-function mutation in Cys_122_ (c.366 A>T; p.Cys122Tyr) (C122Y) with a high prevalence for ventricular arrhythmias, which in the case of the proband required implantation of an intracardiac defibrillator (ICD). To study the molecular mechanisms underlying life-threatening arrhythmias produced by the Kir2.1^C122Y^ mutation, we generated a mouse model of ATS1 using adeno-associated virus (AAVs) Kir2.1^C122Y^ gene transfer that recapitulates the ATS1 phenotype. We used a multidisciplinary approach that included patch-clamping, electrophysiological stimulation, as well as molecular biology, molecular dynamic (MD) modelling, and bioluminescence resonance energy transfer (BRET) to demonstrate that a disulfide bond break in the Kir2.1 extracellular domain disrupts PIP_2_-dependent regulation, leading to channel dysfunction and triggering life-threatening arrhythmias.

## Materials & Methods

### See Supplemental Methods for more detail

#### Ethics Statement

All animal experiment procedures conformed to EU Directive 2010/63EU and Recommendation 2007/526/EC. Skin biopsies were obtained from one patient carrying the Kir2.1 C122Y mutation after written informed consent, and consent to publish, in accordance with the Ethical Committee for Research of CNIC and the Carlos III Institute (CEI PI58_2019-v3), Madrid, Spain. Animal protocols were approved by the local ethics committees and the Animal Protection Area of the Comunidad Autónoma de Madrid (PROEX 111.4/20).

#### Mice

C57BL/6J mice, 4-5-weeks-old, were obtained from the Charles River Laboratories, and reared and housed in accordance with CNIC animal facility guidelines and regulations.

#### Adeno-associated virus vector production, purification and mouse model generation

AAV vectors were generated using the cardiomyocyte-specific cardiac TroponinT proximal promoter (cTnT) and encoding wildtype Kir2.1 (Kir2.1^WT^) or the ATS1 Kir2.1 mutant (Kir2.1^C122Y^), followed by tdTomato report. Vectors were packaged into AAV serotype 9 (AVV9) and produced by the triple transfection method, using HEK293T cells as described previously^20,21^. Mice were anesthetized with ketamine (60 mg/kg) and xylazine (20 mg/kg) via the intraperitoneal (i.p.) route. Thereafter, 3.5×10^10^ virus particles were inoculated intravenously (i.v.) through the femoral vein in a final volume of 50μL. Only well-inoculated animals were included in the studies. All experiments were performed 8-to-10 weeks after infection. *Ex-vivo* fluorescent signal confirming cardiac expression and distribution of protein expression was assessed as described.^22^

#### Echocardiography

Transthoracic echocardiography was performed blindly by an expert operator using a high-frequency ultrasound system (Vevo 2100, VisualSonics Inc., Canada) with a 40-MHz linear probe, and analyzed as described (in *Supplemental Methods)*.

#### Surface ECG recording

Mice were anesthetized using isoflurane inhalation (0.8-1.0% volume in oxygen). Four-lead surface ECGs were recorded for 5 minutes using subcutaneous limb electrodes connected to an MP36R amplifier unit (BIOPAC Systems). Data acquisition and analysis were performed using AcqKnowledge software.

#### *In-vivo* intracardiac recording and stimulation

An octopolar catheter (Science) was inserted through the jugular vein and advanced into the right atrium (RA) and ventricle (RV) as previously described.^23^ Atrial and ventricular arrhythmia inducibility was assessed by applying consecutive trains at 10Hz and 25Hz, respectively.

#### Cardiomyocyte isolation

The procedure was performed as described by Macías et al.^24^

(See *Supplemental Methods)*.

#### Membrane fractionation, immunoprecipitation and immunoblotting

Total protein was obtained from isolated Kir2.1^WT^ and Kir2.1^C122Y^ cardiomyocytes using RIPA buffer (150 mM NaCl, 10 mM Tris-HCl, 1 mM EDTA, 1% Triton X-100, 0.1% SDS and 0.1% Sodium deoxycholate) supplemented with protease inhibitor cocktail (Roche) and quantified by BCA protein assay (Bio-Rad). A total amount of 50 µg of protein was resolved in each lane on 10% SDS-PAGE gels, electrotransferred onto 0.2 µm PVDF membrane (BioRad) and probed with specific antibodies. For membrane fractionation, cells were extracted and homogenized in ice-cold homogenization medium. After lysis, protein extract was processed according to manufacturer’s specifications (Abcam). See further details in the *Supplemental Methods*,

#### Bioluminescence Resonance Energy Transfer (BRET) Lipid binding assay

HEK293T cells were transfected with 2 µg of plasmid encoding Kir2.1^WT^ or Kir2.1^C122Y^ protein fused with Nluc (nanoluciferase) in the C-terminal region. After 48h, the BRET assay was done in a 96-well plate as previously described.^25^ See *Supplemental Methods* for details.

#### Patch-clamping in isolated cardiomyocytes

The whole-cell patch-clamp technique and data analysis procedures and internal and external solutions (***Supplementary Table 1***) were similar to those previously described.^9–13^ Details are presented in the *Supplemental Methods*.

#### Calcium dynamics assays

Cytosolic Ca^2+^ was monitored according to previously described protocols.^26–28^ Briefly, cells were loaded with Fluo-4-AM (Invitrogen). Fluorescence was detected in line scan mode (usually 2 ms/scan), with the line drawn approximately through the center of the cell and parallel to its long axis.

#### Dynamic modeling to predict Kir2.1-PIP_2_ interaction

For each monomer we used the pre-opened state of Kir2.2 bound to PiP_2_ as a template (PDB code 3SPH) to conduct molecular dynamics (MD) modelling. We generated homology PiP_2_ models binding to Kir2.1^WT^, Kir2.1^C122Y^ homotetramer and Kir2.1^WT/C122Y^ heterotetramer to study Kir2.1-PiP_2_ interactions using 2000 ns MD. The CHARMM-GUI server allowed us to simulate both membrane and environment. - Please see *Supplemental Methods* for detailed description of the procedures.

#### Statistical analyses

We used GraphPad Prism software version 7.0 and 8.0. In general, comparisons were made using Student’s t-test. Unless otherwise stated, we used one- or two-way ANOVA for comparison among more than two groups and Tukey correction for multiple comparisons. Data are expressed as mean ± SEM, and differences are considered significant at p<0.05.

## Results

### Life-threatening arrhythmias in an ATS1 family with the Kir2.1^C122Y^ mutation

We screened a family with members suffering numerous idiopathic sudden loss-of-consciousness episodes using a targeted sequencing gene panel involved in arrhythmias (*RYR2, CASQ2, TRDN, CALM1 and KCNJ2*). We identified a novel *de novo* potential pathogenic heterozygous missense variant c.365 A>T; p.Cys122Trp of the *KCNJ2* gene for ATS1 (LQTS type 7) in two family members (**Figure 1A-B**). The proband (patient II.2) was a 16-year-old female of Caucasian origin who experienced several sudden loss of consciousness events of unknown origin. Initially, patient II.2 was diagnosed with mitral valve prolapse of the anterior leaflet without hemodynamic repercussions. The electrophysiological study was negative following a hospital admission for syncope and subsequent evidence of polymorphic ventricular extrasystoles refractory to antiarrhythmic drugs (propafenone, mexiletine and lidocaine). She continued with propranolol treatment (120 mg/d) combined with oral mexiletine (200mg/8h). At age 23, a single-chamber cardioverter-defibrillator (ICD) was implanted after several episodes of syncopal polymorphic ventricular tachycardia (PVT) and registering three appropriate discharges throughout ages 25-35 during sodium channel blocker administration (**Figure 1C**). ECG analysis revealed a corrected QT (QTc) interval in the upper limit of the normal range (470 ms) with pronounced U waves and polymorphic extrasystoles with frequent trigeminy episodes (**Figure 1D**). She is now 45 and currently under treatment with nadolol 120 mg/d and spironolactone 25 mg/d. The proband’s son (patient III.1) remains asymptomatic at the age of 8. However, ECG analysis revealed a prolonged QTc interval of 490 ms with a widened T wave and prominent U waves (**Figure 1E**), consistent with ATS1 symptoms.

**Figure 1.**
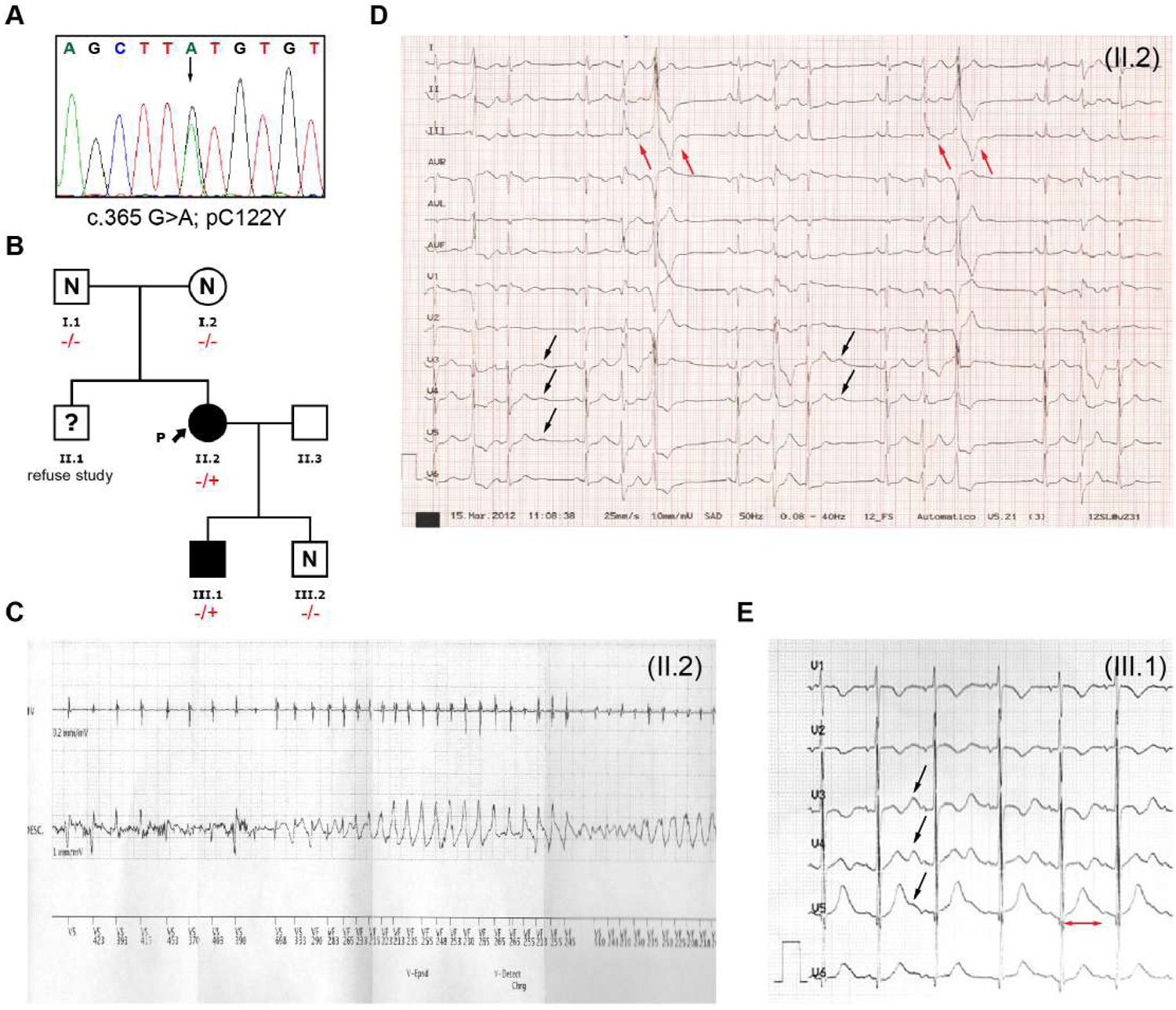
Genetics and ECG phenotype of ATS1 family members with Kir2.1^C122Y^ mutation. **A**: DNA sequences derived from proband’s genomic DNA. The trace shows a heterozygous substitution of guanine to adenine resulting in the C122Y amino acid change. **B**: Family pedigree according to the carrier status of the p.Cys122Tyr *KCNJ2* gene variant. Males and females are marked with squares and circles, respectively. Mutation carriers are marked with (−/+) and non-carriers with (-/-). Uncertain mutation carriers are marked with (?) and non-affected with (N). Phenotype positive individuals are marked in black. Proband is indicated with black arrow and (P). **C.** Twelve-lead ECG of proband (II.2) at age 23, showing an episode of syncopal polymorphic ventricular tachycardia during sodium channel blocker (Mexiletine; 600mg/day) treatment combined with β-blocker therapy (Propranolol; 20mg/12h) **D**: ECG of the proband (II.2) showing typical Andersen-Tawil Syndrome abnormalities. Prominent U waves are marked by black arrows. Bidirectional ventricular extrasystoles are marked by red arrows. **E**: ECG from individual III.1 demonstrating genotype-phenotype segregation. Prominent U waves are marked with black arrows. Broad T wave and QTc interval prolongation is marked by a red line (510 ms).

### Cardiac conduction defects and arrhythmias in Kir2.1^C122Y^ mice

We used intravenous AAV-mediated cardiac specific gene transfer^22^ to generate mice expressing Kir2.1^WT^ or Kir2.1^C122Y^. We confirmed AAV infection throughout the heart and that cardiomyocytes stably expressed the specific targeted transgenes (**Supplemental Figure 1**), with no cardiac morphological changes or contractile dysfunction evaluated by echocardiography **(Supplemental Figure 1C and Sup. Figure 2).** On surface ECG, Kir2.1^C122Y^ mice showed conduction alterations characteristic of the disease (**Figure 2A**). More importantly, Kir2.1^C122Y^ mice had frequent premature ventricular complexes (PVCs) and runs of non-sustained PVT (**Figure 2B**) in agreement with the ATS1 patient’s phenotype. Under stress conditions induced by isoproterenol (ISO, 5mg/Kg), Kir2.1^C122Y^ mice developed PR and QRS prolongation. Compared with control, Kir2.1^C122Y^ animals exhibited repolarization abnormalities with prolongation of the QT interval and occasional overlap of the T wave with the P wave of the following complex (**Figure 2C-D**). Intracardiac stimulation of the right atrium or ventricle used consecutive trains of stimuli at 10 and 25 Hz. Under basal conditions, Kir2.1^C122Y^ mice had a significantly increased arrhythmia susceptibility with respect to Kir2.1^WT^ (**Figure 2E-F)**; upon stimulation, 5 out of 9 Kir2.1^C122Y^ mice (55,5%) developed atrial or ventricular arrhythmias, including PVT, compared to 0 out of 7 Kir2.1^WT^ mice (0%). ISO administration increased arrhythmia susceptibility in both atria and ventricles of Kir2.1^C122Y^ (8 out of 9 mice, 88,9%), vs. Kir2.1^WT^ (1 out of 7 mice, 14,2%). Altogether, these results indicate that the Kir2.1^C122Y^ mutation recapitulates the ATS1 patient’s cardiac electrical phenotype, establishing an arrhythmogenic substrate.

**Figure 2.**
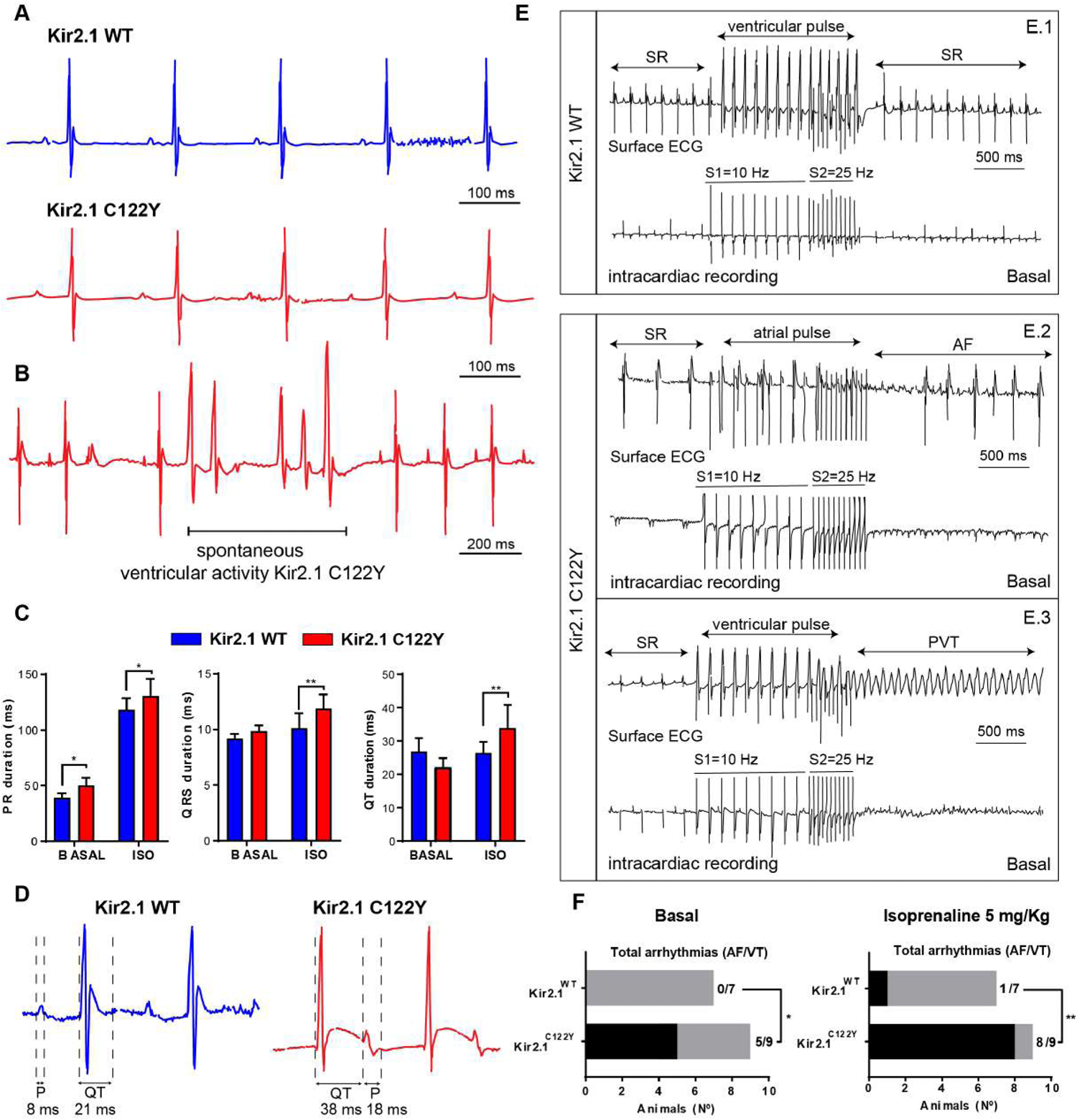
Kir2.1^C122Y^ mice recapitulate the ATS1 patientś phenotype and increased susceptibility to arrhythmias. **A**: Representative lead-II ECG recordings from AAV-transduced Kir2.1^WT^ (top) and Kir2.1^C122Y^ (bottom) mice. The record shows normal sinus rhythm with prolonged PR interval in mutant animals (N= 7 animals per group). **B**: ECG in a Kir2.1^C122Y^ animal showing frequent premature ventricular complexes (PVCs) manifested as duplets. **C-D**: Effects of isoprenaline (ISO, 5 mg/Kg) administration on electrical conduction and QT interval in Kir2.1^C122Y^ animals compared to basal condition (N= 7 animals per group). **E**: Representative lead-II ECG traces (top) and corresponding intracardiac recordings (bottom) before (SR; sinus rhythm), during and after intracardiac application of stimulus trains at 10 and 25 Hz under basal conditions. **E.1**, atrial stimulation in a Kir2.1^WT^ mouse failed to induce an arrhythmia. **E.2**, atrial stimulation in a Kir2.1^C122Y^ mouse induced a period atrial fibrillation. **E.3**, ventricular stimulation in a Kir2.1^C122Y^ mouse induced polymorphic ventricular tachycardia (PVT). **F**: Contingency plots of number of animals with arrhythmogenic response after intracardiac stimulation at baseline, and after treatment with ISO (5 mg/Kg). Each value is the mean ± SEM (N=7-9 animals per group). Statistical analysis by two-tailed ANOVA and Student-t test. * p<0.05; ** p<0.01.

### Kir2.1^C122Y^ subunits are able to form heterotetramers

Kir2.1 channels can exist either as homo- or hetero-tetrameric complexes consisting of either four identical Kir2.1 subunits or in various combinations with the structurally related members of the Kir2.x subfamily of inward rectifier K^+^ channels^29^. To clarify the mechanisms by which the C122Y mutation causes channel dysfunction in ATS1, we determined whether Kir2.1^C122Y^ can assemble with WT subunits and traffic to the surface membrane (**Supplemental Figure 3**). Immunoprecipitation studies using differently tagged Kir2.1 subunits were used to test whether the mutation affected subunit assembly. The HA and Myc epitope tags were incorporated into an external site that does not perturb channel activity^30,31^ (**Supplemental Figure 3A**). In these studies, HEK293T cells were either co-transfected with Myc-tagged Kir2.1 (WT) or HA-tagged Kir2.1 (WT or C122Y) at a 1:1 ratio. Recovered immunoprecipitants on anti HA-bound beads were resolved by SDS-PAGE, and the extent of HA-tagged channel subunit interaction was assessed using anti-Myc antibodies in immunoblots. As shown by the representative experiment (**Supplemental Figure 3B**), the wild-type Myc-Kir2.1 co-immunoprecipitated with both HA-tagged subunits, indicating the mutation does not alter subunit interaction. In addition, immunocytochemical analysis of co-transfected cells revealed that the Myc-tagged Kir2.1^WT^ and HA-tagged Kir2.1^C122Y^ subunits are highly co-localized (**Supplemental Figure 3C**), offering further evidence that the C122Y subunits are capable of assembling with the WT subunits in cells.

### Kir2.1^C122Y^ subunits traffic to the cardiomyocyte surface membrane

Kir2.1 localizes at two separate well-defined striated microdomains running parallel to each other at ∼0.9 µm intervals throughout the cardiomyocyte.^24^ One microdomain corresponds with the t-tubules where Kir2.1 co-localizes with the voltage gated cardiac sodium channel Na_V_1.5 (∼1.8 µm spacing). The other is at the sarcoplasmic reticulum (SR) where Kir2.1 functions to control calcium homeostasis (**Figure 3A** and **B**).^24^ Disruption of one or both microdomains leads to malfunction of Kir2.1 and Na_V_1.5 channels that might trigger arrhythmias. However, unlike the defective distribution pattern that was demonstrated for the trafficking deficient mutation Kir2.1^Δ314–315,24^ immunolocalization and confocal image analysis of isolated ventricular cardiomyocytes from Kir2.1^C122Y^ animals revealed an unaltered distribution pattern for both Kir2.1 and Na_V_1.5 channels (**Figure 3A** and **B**). When we determined the percentage of membrane expression using an anti-Na^+^/K^+^ ATPase immunostaining, the results again showed a similar distribution of Kir2.1 and Na_V_1.5 channels in Kir2.1^WT^ and Kir2.1^C122Y^ cells, with a small but significant reduction in Na_V_1.5 accumulation level in mutant cardiomyocytes (**Figure 3C**). Similarly, on western blot, Na_V_1.5 protein expression was lower for the mutant cardiomyocytes with a trend toward a decrease in total protein (**Figure 3D-E**). Trafficking of both Kir2.1 and Na_V_1.5 to their membrane microdomains depends in part on their classical route that involves incorporation into clathrin-coated vesicles at the trans-Golgi network marked by interaction with the adaptor protein complex-1 ϒ-adaptin subunit (AP-1).^32^ Trafficking may also occur via an unconventional route directly from SR in a GRASP dependent manner.^33^ To test whether Kir2.1^C122Y^ disrupts Kir2.1 trafficking we analyzed AP-1 and GRASP65 proteins by immunofluorescence of both WT and mutant cardiomyocytes. As shown in **Figure 3F-G**, the AP-1 expression profile was identical in both groups. Similarly, GRASP65 staining presented an F-function distribution (distance from particles to nearest neighbor particle) with no differences in either WT or mutant groups. Also, co-localization of Kir2.1 with GRASP65 was similar in both groups. From the foregoing the Kir2.1^C122Y^ variant is able to form heterotetramers with WT subunits and retains trafficking ability. Taken together, these observations strongly suggest that the C122Y mutation leads to cardiac electrical alterations via mechanisms other than those recently demonstrated for the trafficking deficient Δ314-315 mutation.^36^ This led us to further explore the biophysical and electrophysiological properties of the Kir2.1^C122Y^ channel and determine whether the mutation directly alters potassium conductance and/or disrupts protein stability.

**Figure 3.**
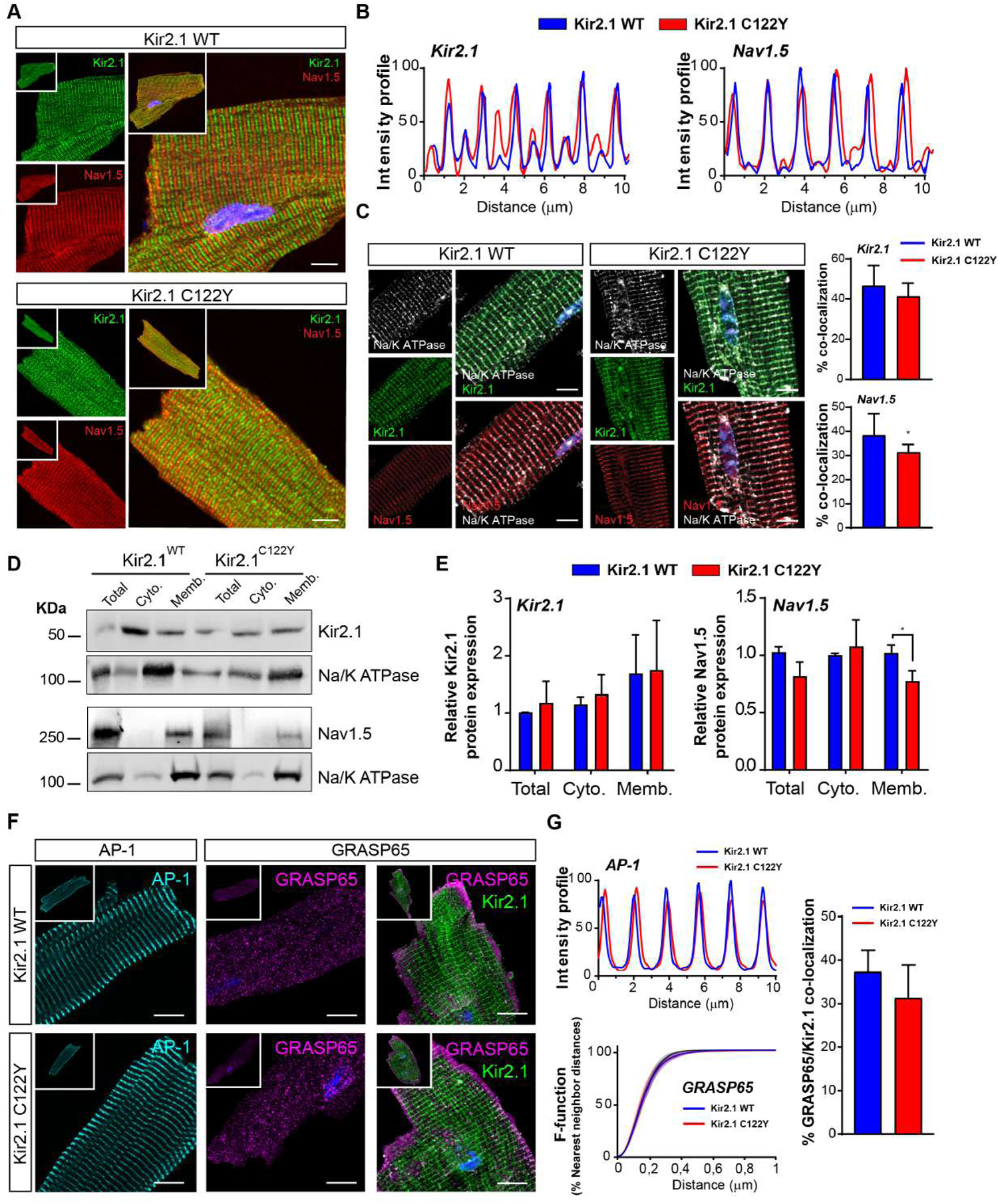
Kir2.1^C122Y^ cardiomyocytes preserve Kir2.1 and Na_V_1.5 protein trafficking, but both proteins are reduced at the sarcolemma. **A**: Confocal images of Kir2.1 and Nav1.5 channels in Kir2.1^WT^ and Kir2.1^C122Y^ cardiomyocytes. Scale bar, 10μm. **B**: Fluorescence intensity profiles show distribution patterns for both Kir2.1 (left panel) and Na_V_1.5 (right panel) channels in WT and Kir2.1^C122Y^ cardiomyocytes. Note double banding for Kir2.1 indicating SR expression.^24^ **C**: Representative immunofluorescence images show co-localization of Kir2.1 (green) and Na_V_1.5 (red) with Na^+^/K^+^ ATPase (white) at the sarcolemma. Graphs show percentage of co-localization with significantly reduced Na_V_1.5 (* p<0.05; t test). Scale bar, 10μm **D**: Western blots comparing cytosolic and sarcolemmal Kir2.1 and Na_V_1.5 in Kir2.1^WT^ vs Kir2.1^C122Y^ cardiomyocytes. Data were normalized using Na^+^/K^+^ ATPase. **E**: Graphs show western blot quantification of cytosolic and sarcolemmal Kir2.1 and Na_V_1.5 channels. Note reduced Na_V_1.5 at the sarcolemma (N=4-5 animals per group) (* p<0.05; two-tailed ANOVA). **F**: Confocal images of classical (AP-1) and unconventional (GRASP65) trafficking routes for Kir2.1 and Na_V_1.5. Scale bar, 10μm. **G**: Quantification of fluorescence intensity profiles for AP-1, F-function (% nearest neighbour distances) and percentage of GRASP co-localization in isolated Kir2.1^WT^ and Kir2.1^C122Y^ cardiomyocytes. (N=3 animals per group; n=7-9 cells). (* p<0.05; two-tailed ANOVA). Scale bar, 10μm. Each value is the mean ± SEM.

### Kir2.1^C122Y^ cardiomyocytes exhibit defects in excitability and action potential duration

We performed patch-clamping experiments in isolated cardiomyocytes from Kir2.1^WT^ and Kir2.1^C122Y^ expressing hearts. We focused on both I_K1_ and the sodium inward current (I_Na_) to test whether the impulse conduction disturbances and arrhythmias observed in this model of ATS1 are due to defects in one or both currents. The results show a 90% reduction in the outward I_K1_ density of Kir2.1^C122Y^ compared with Kir2.1^WT^cardiomyocytes (**Figure 4A**), which explains why we were unable to obtain reliable current clamp measurements of AP characteristics, since the vast majority of Kir2.1^C122Y^ cardiomyocytes (11 out of 12 or 91.7%) tested were substantially depolarized at rest (∼-35mV) and unable to generate APs upon stimulation. Furthermore, surprisingly, Kir2.1^C122Y^ cardiomyocytes showed a slight but significant decrease in I_Na_ density compared with controls (**Figure 4B**) with no significant changes in the voltage-dependence of current activation or inactivation (**Supplemental Figure 4**). These data further demonstrate that, while the mutant Kir2.1 protein traffics and is expressed at the membrane, it is dysfunctional and also reduces Na_V_1.5 function. Altogether, these results reinforce the hypothesis that conduction disturbances and arrhythmias in ATS1 patients are due to a defect in cardiomyocyte excitability.

**Figure 4.**
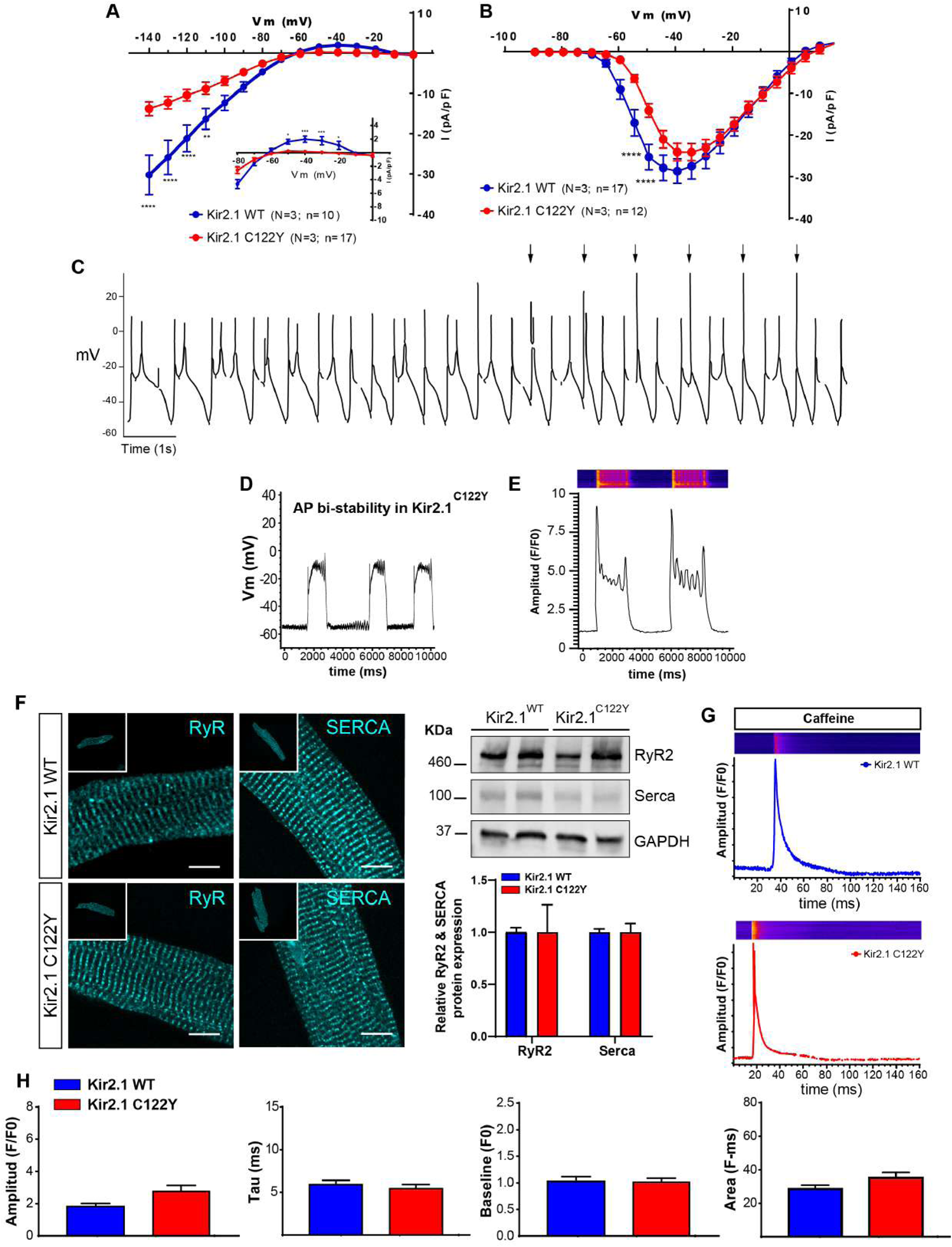
Kir2.1^C122Y^ alters electrophysiology in isolated mouse cardiomyocytes. **A**: Superimposed I_K1_ current-voltage (IV) relationships for Kir2.1^WT^ (blue) and Kir2.1^C122Y^ (red) cardiomyocytes. **B**: Superimposed I_Na_ IV relationships for Kir2.1^WT^ (blue) and Kir2.1^C122Y^ (red) cardiomyocytes. **C**: Representative action potential time series recorded during current-clamping in an isolated Kir2.1^C122Y^ cardiomyocyte. Note spontaneous action potentials with excessively long APD generating early afterdepolarizations (EADs) and triggered activity. **D**: Membrane potential bi-stability in a Kir2.1^C122Y^ mutant with EADs appearing above −20 mV. Graph shows quantification of bi-stability events in a Kir2.1^C122Y^cardiomyocyte. **E:** Representative confocal image and profile of calcium transient dynamics in another isolated Kir2.1^C122Y^ cardiomyocyte. Note amplitude bi-stability and large numbers of spontaneous calcium release events spreading throughout the cell. **F**: Left, Immunolocalization of ryanodine receptor (RyR_2_) and Ca^2+^-ATPase (SERCA) in AAV-transduced ventricular cardiomyocytes from Kir2.1^WT^ and Kir2.1^C122Y^ mice. Scale bar, 10μm (N=3 animals per group; n=7-8 cells). Right, western blots showing similar amounts of total protein for both (N=4 animals per group). **G**: Representative fluorescence profiles of caffeine-induce calcium release in Kir2.1^WT^ and Kir2.1^C122Y^ cardiomyocytes. **H:** Graphs show amplitude, Tau (Decay kinetics) and Baseline of each Ca^2+^ transient, as well as the total area) (N=3 animals per group; n=10-17 cells). Each value is represented as the mean ± SEM. Statistical analyses were conducted using two-tailed ANOVA. * p<0.05; ** p<0.01; **** p<0.0001.

As illustrated in **Figure 4C**, AP recordings revealed that, the 7 out of 12 (58,3%) Kir2.1^C122Y^ cardiomyocytes that remained excitable after isolation generated significantly prolonged APs, early afterdepolarizations (EADs), triggered discharges and bi-stability of the RMP (**Figure 4D**). Accordingly, we analyzed the intracellular calcium dynamics in both WT and ATS1 mice. Confocal images of Ca^2+^ dynamics showed that Kir2.1^C122Y^ cardiomyocytes had an excitation-contraction (e-c) coupling defect with multiple abnormal spontaneous calcium release events during systole and diastole (**Figure 4E**). Since Ca^2+^ movements across the sarcoplasmic reticulum (SR) are controlled by the ryanodine receptor (RyR_2_)-mediated Ca^2+^ release and the Ca^2+^-ATPase (SERCA)-mediated Ca^2+^ reuptake to-and-from the cytosol and SR lumen, we wondered whether protein alteration could happen in the Kir2.1^C122Y^ mouse model. However, confocal images of protein localization profiles were identical in Kir2.1^C122Y^ and Kir2.1^WT^ cardiomyocytes, and total protein levels were also similar (**Figure 4F**). Since, K^+^ flux across Kir2.1 SR channels contributes countercurrent to Ca^2+^ movement, ^24^ we analyzed the intracellular Ca^2+^ dynamics in both controls and ATS1 mice (**Figure 4G-H**). Cardiomyocytes expressing Kir2.1^WT^ and Kir2.1^C122Y^ showed similar Ca^2+^ transient decay under acute caffeine administration in intact cardiomyocytes (**Figure 4G-H**). These results indicate that the Ca^2+^ alterations are due to functional defects at the sarcolemma, including RMP depolarization and reduced excitability, rather than Kir2.1 dysfunction at the SR.

### Disulfide bond loss reorganizes tridimensional channel structure interfering with Kir2.1^C122Y^-PIP_2_ binding

Cys_122_ localizes at the extracellular loop of the Kir2.1 channel, immediately after the first transmembrane domain, where it is cross-linked by an intramolecular disulfide bond with Cys_154_ at the beginning of the second transmembrane α-helix (**Figure 5A**). Both residues and their disulfide bond are conserved across the Kir family (**Figure 5B**), which is crucial for proper channel folding, as they may help accommodate the extracellular loop in an optimal tridimensional structure.^18^ We used *in-silico* homology modelling to derive predictions of the molecular structure of the Kir2.1^C122Y^ mutant channel, and thus understand the possible mechanisms underlying its dysfunction. Atomic level modelling showed that, compared to the WT channel, Kir2.1^C122Y^ undergoes a clear reorganization (TMscore 0.73; RDMS: ∼6Å for homotetramer or TMscore 0.78; RDMS: ∼7Å for heterotetramer) (**Figure 5C-D**). The Gibbs free-energy values for Kir2.1^C122Y^ were more positive compared to WT (WT: 4801.404 vs C122Y: −4131.754 for homo or −2274.207 for heterotetramer) (**Figure 5D**). This indicates a more unstable state in Kir2.1^C122Y^ homo- and heterotetrameric channels, suggesting that the incorporation of mutant subunit could affect the integrity of the WT monomers or even affect the macromolecular channelosome complex, including Kir2.1 and Na_V_1.5.

**Figure 5.**
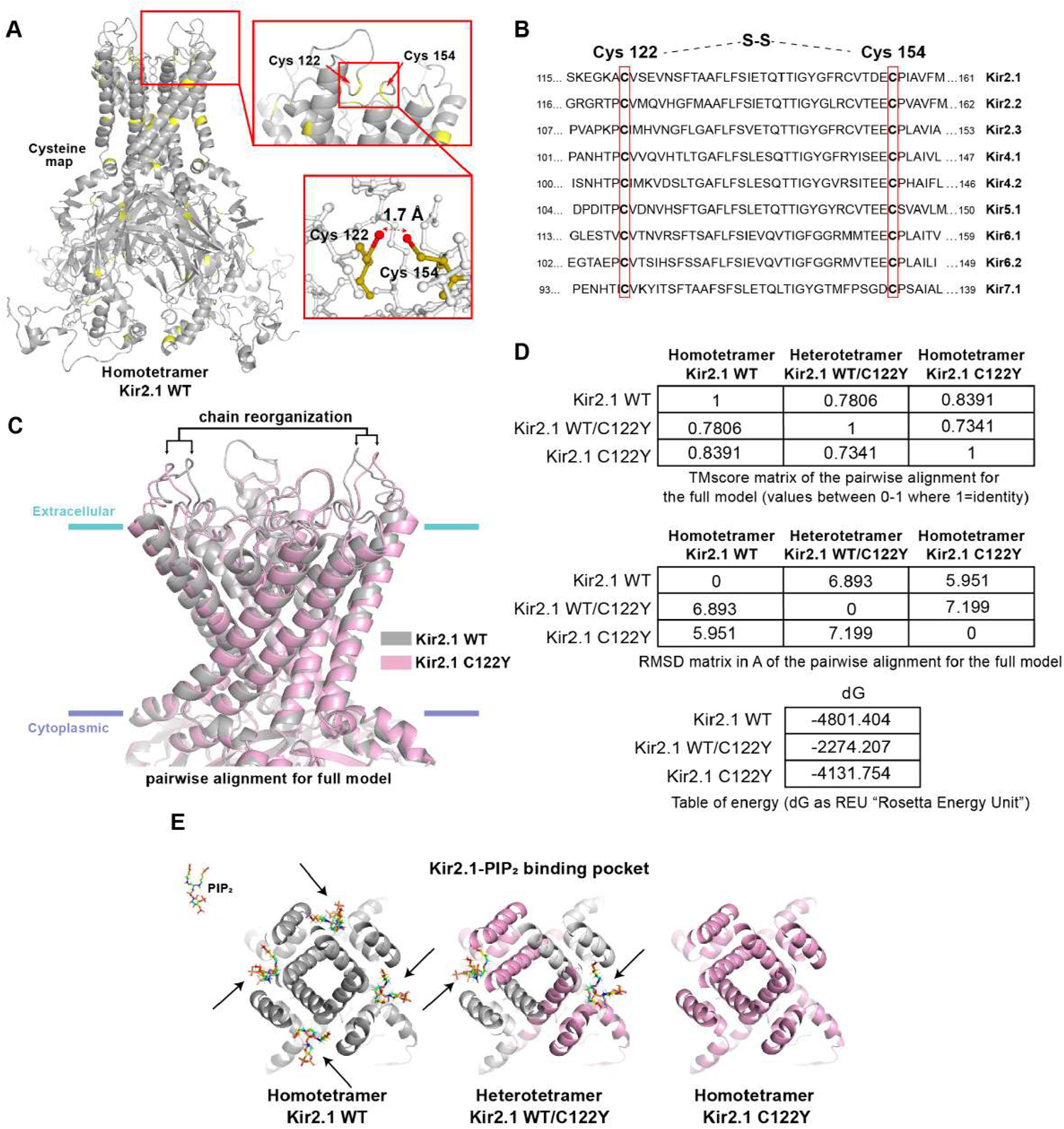
The C122Y mutation alters Kir2.1 channel conformation and PIP_2_ binding. **A**: Topological scheme of Kir2.1 homotetramer channel indicating cysteine positions (yellow). **B**: Amino acid sequence in Kir family indicating highly conserved extracellular disulfide bond. Cys122 and Cys154 are indicated in Kir2.1 **C**: Pairwise alignment for full model (Grey, Kir2.1^WT^; pink, Kir2.1^C122Y^). **D**: Upper panel, TMscore matrix of the pairwise alignment for the full model. Values between 0-1, where 1 is the identity. RMSD matrix (middle panel) in angstroms (Å). Lower panel, Table of Gibbs free-energy values (dG) of WT and mutant homo- and heterotetramer. **E**: Docking modelling of Kir2.1-PiP_2_ interaction in Kir2.1^WT^, homo- and heterotetramers of Kir2.1^C122Y^(see text for detailed explanation of each panel).

To predict Kir2.1-PIP_2_ interaction ability we incorporated PIP_2_ molecules in the simulation. Our results showed an altered Kir2.1^C122Y^-PIP_2_ interaction following a dominant-negative pattern. The Kir2.1^WT/C122Y^ heterotetramer presented 2 out of 4 PIP_2_ molecules compared with the complete set of 4 PIP_2_ in the Kir2.1^WT^ homotetramer, one per monomer (**Figure 5E**). Notably, the Kir2.1^C122Y^ homotetramer abolished completely PIP_2_ interaction, in accordance with the I_K1_ current suppression in homozygous mutant conditions in C154F^34^ and C122Y-expressing HEK cells (**Supplemental Figure 5**). Taken together, these *in-silico* homology experiments predict that the loss of the highly-conserved extracellular Cys122-to-Cys154 disulfide bond in channels containing the Kir2.1^C122Y^ isoform may result in a clear atomic re-structuration with loss of function by mechanisms that include, at least in part, a pronounced interference with the PIP_2_ binding pocket.

### Cys122-Cys154 disulfide bond breakup disrupts Kir2.1-PiP_2_ interaction dynamics

We conducted *in-silico* molecular dynamics (MD) studies to more rigorously establish whether the extracellular Cys122-to-Cys154 disulfide bond breakup in the Kir2.1^C122Y^ mutant channel disrupts Kir2.1-PIP_2_ interaction (**Figure 6**). We generated Kir2.1 homology models bound to a single PIP_2_ molecule per monomer in Kir2.1^WT^, Kir2.1^C122Y^ homotetramer and Kir2.1^WT/C122Y^ heterotetramer to study Kir2.1-PIP_2_ interactions throughout an individual 2000 ns MD replica (see *Supplemental Methods* for details of the overall approach). For each monomer we used the pre-opened state of Kir2.2 bound to PIP_2_ as a template (PDB code 3SPH).^35^ The CHARMM-GUI server allowed us to simulate both membrane and environment (**Figure 6A-B**); we then performed three independent replicas for each model. First, we evaluated the conformational changes in the extracellular space by monitoring either C_122_ or Y_122_ backbone dihedral angles along the 2000 ns MD. Comparative analysis showed only 28% conserved-frames in backbone dihedral angles, while 72% presented a shift in Φ-dihedral angle from around −70° to −140° shortly after the first 100 ns (**Figure 6C and Supplemental Figure 6**). The Y_122_ sidechain reorientation resulted in movement of D_112_, and consequent break of the internal hydrogen-bonding network between D_112_ and the H_110_ sidechain and the NH backbone of C_122_ within the extracellular loop (**Supplemental Figure 7**). Thus, hydrogen bonds between the H_110_ sidechain and the Y_122_ backbone were either absent or generally present in less than 50% of the frames. In addition, in several of the MD simulations a new hydrogen bond was formed between D_112_ and K_117_. Therefore, C122Y leads to a reorganization of the hydrogen-bonding network of the extracellular loop that might alter Kir2.1 function. (**Supplemental Table 2**). Nevertheless, neither of the two Y_122_ conformations observed in the MD led to a significant change in the relative disposition of the outer and inner helices, as shown by the measurement of the distance between two opposite residues (I_106_ and I_156_) located near the extracellular side of each of those helices (**Supplemental Figure 8**).

**Figure 6.**
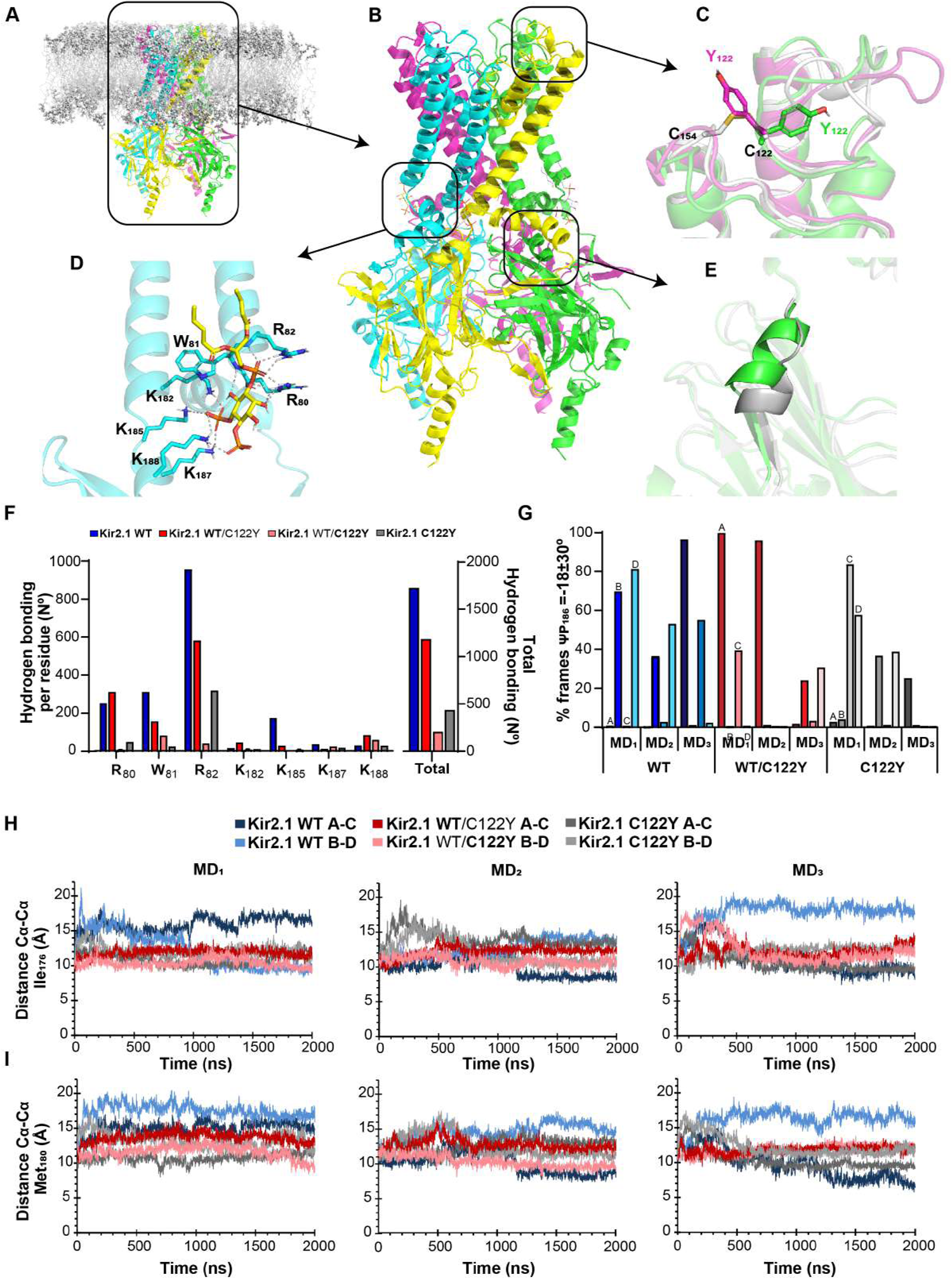
Extracellular disulfide bond break reduces PiP_2-_dependent Kir2.1 regulation. **A**: Schematic representation of Kir2.1 tetramer embedded in a bilipid layer. **B**: Structure of Kir tetramer. Monomers are represented in different colors. **C**: Illustrative C122 or Y122 sidechain orientation. Superposition of Kir2.1^WT^ (grey) and two representative Kir2.1^C122Y^ monomers (in green the most frequent Y122 orientation, in purple, the minor one). **D**: Representative illustration of hydrogen bond network between Kir2.2 and PIP_2_. Same hydrogen bondings as in the generated homology model were tested for Kir2.1. **E**: Evolution of the C-linker during the MD: from a helix (green) to a less structured linker, as shown by a representative 2000 ns snapshot (grey). **F**: Histogram representing the average number of PIP_2_-Kir2.1 hydrogen bonds per residue along the 2000 ns simulation, for Kir2.1^WT^ (blue), Kir2.1^WT/C122Y^ (grey) and Kir2.1^WT/C122Y^ (red). These values are the average of the three replicas and the four chains for each tetramer. **G**: Histogram representing the percentage of frames in which the ψ dihedral angle of the Pro186 is within those expected for a 3_10_ helix (ψ =-18±30°). For Kir2.1^WT/C122Y^, A and C represent the non-mutated monomers. **H**: I_176_ and M_180_ Cα-Cα distances between two opposite monomers along the 2000 ns MD. Color code on top. N=3 replicates.

Kir2.1-PIP_2_ interactions involve hydrophobic contacts with PIP_2_ acyl chains, and more specific polar interactions between PIP_2_ phosphates and positively charged Kir2.1 residues at the transmembrane domain (TMD)-to-cytoplasmic domain (CTD) interface.^35,36^ A detailed study of the atomic Kir2.1-PIP_2_ hydrogen-bonding distance yielded a global loss of hydrogen bonds in the Kir2.1^C122Y^ channels that directly affected PIP_2_ interactions. Comprehensive analysis of the R_80_W_81_R_82_ motif and the lysine-cluster K_182_K_185_K_187_K_188_ of the helicoidal CTD-to-TMD linker (C-linker) (**Figure 6D**) showed a clear reduction in hydrogen bonding capacity in both hetero- and homotetrameric Kir2.1^C122Y^ channels throughout the 2000 ns MD (**Figure 6F and Supplementary Table 3**). Our simulations predict that, compared with the Kir2.1^WT^ tetramers, Kir2.1^WT/C122Y^ and Kir2.1^C122Y^ channels progressively lose the characteristic hydrogen bond of the PIP_2_ 1′ phosphate with R_80_W_81_R_82_, particularly R_82_, and the PIP_2_ 4′ and 5′ phosphates with the C-linker (**Supplemental Table 3**). As expected, the unmutated chains (chains A and C) in Kir2.1^WT/C122Y^ heterotetramers showed a similar behavior to WT chains (**Supplemental Table 3**).

Upon PIP_2_ binding at the interface between TMD and CTD, the C-linker undergoes a disorder-to-order transition bringing both domains closer together. Thus, the G-loop wedges into the TMD causing the inner helix gate to open.^35^ Follow-up of this transition showed that the C-linker loses the hydrogen bonds characteristic of α-helix structures in the Kir2.1^C122Y^ hetero- and homotetramer. In comparison, Kir2.1^WT^ maintained a more pronounced hydrogen-bond network between K_188_-E_191_ and R_189_-T_192_, as well as the K_185_, P_186_, K_187_ and N_190_ residues, indicating that in the mutant channels the N- and C-terminal of the C-linker helix were destructured faster than WT (**Figure 6E and Supplemental Table 4**). Interestingly, at the beginning of the C-linker motif, the dihedral angles of P_186_ were within those of 3_10_ helix φ (−71) and ψ (−18) in the PIP_2_-bound Kir2.1^WT^ structures. However, in the Kir2.1^C122Y^ hetero- and homotetramer the φ dihedral varied in correlation with a progressive loss of the C-linkeŕs helical character (**Figure 6G**). Compared with Kir2.1^WT^, the Kir2.1^C122Y^ homotetramer had a lower percentage of frames corresponding to the dihedral 3_10_ helix in P_186_ (Kir2.1^C122Y^: 17% at 1000 ns, 8% at 2000 ns; Kir2.1^WT^: 33% at 1000 ns, 17% at 2000 ns), the percentages of the heterotetramer being intermediate (**Figure 6G**).

Finally, we measured the distance between Cα carbons of representative pore constriction Ile_176_ and Met_180_ residues at the TM and A_306_ of the G-loop from opposite chains to study the pore opening state during the 2000 ns MD (**Figure 6H and Supplemental Figure 8**)^36,37^. For both Ile_176_ and Met_180_ the distance between the A-B and between the B-D chains decreased progressively in hetero and more pronouncedly in homo mutant channels, with larger values for WT chains in the first 500 ns, which likely correlated with a more open state in WT channels (**Figure 6I**). Longer MD times using WT channels showed that the distance among the Cα carbons of the above residues decreased in two opposite monomers and led to an increase in the distance between the other two monomers, as observed in the gating mechanism for KirBac3.1.^38^ Similarly, the Cα-carbon distance between A_306_ residues in the G-loop decreased in the mutant hetero- and more pronouncedly in the homotetramer (**Supplemental Figure 9**). These results strongly suggest that the extracellular disulfide bond break of Kir2.1^C122Y^ closes the channel by altering the Kir2.1-PIP_2_ hydrogen-bond network, which in the WT stabilizes PIP_2_ function to maintain the open state of the channel.

### Kir2.1^C122Y^ has a reduced sensitivity to, and binding capacity for PIP_2_

To test for PIP_2_ binding to Kir2.1, we fused a nanoluciferase (Nluc) to the C-terminus of the channel and used a soluble fluorescent PIP_2_ (FL-PIP_2_) analog suitable for binding to Kir2.1.^25^ Activation of Nluc produced a FL-PIP_2_-dependent bioluminescence resonance energy transfer (BRET) signal specific for Kir2.1 as shown by the cartoon of the assay design in **Figure 7A**. HEK293T cells were transfected with the WT and C122Y mutant version, respectively, and a bioluminescence assay was performed. We included another Kir2.1 mutant version with a known mutation interfering with PIP_2_-Kir2.1 channel interaction as a negative control (Kir2.1^R218W^). Our results showed that PIP_2_ binds with high affinity to Kir2.1^WT^ but, as expected cannot directly bind to Kir2.1^C122Y^, such as we observed for Kir2.1^R218W^ (**Figure 7B**). To test the sensitivity of Kir2.1 to PIP_2_, we performed inside-out patch-clamping of the Kir2.1^WT^ and the heterozygous condition Kir2.1^WT/C122Y^ currents in co-transfected HEK293T cells at a 1:1 ratio. We recorded I_K1_ in both basal condition and under increasing concentrations of PIP_2_ (25 and 50 μg/ml PIP_2_). The results showed that while in Kir2.1^WT^, PIP_2_ increased the inward K^+^ current in a dose-dependent manner, the Kir2.1^WT/C122Y^ mutation blunted the sensitivity to PIP_2_ (**Figure 7C-D**). Both groups showed an unaltered outward current. Taken together, these results confirm the inability of Kir2.1^C122Y^ channels to functionally interact with PIP_2_ molecules that allow proper channel function, according to the dominant negative effect expected from patient data.

**Figure 7.**
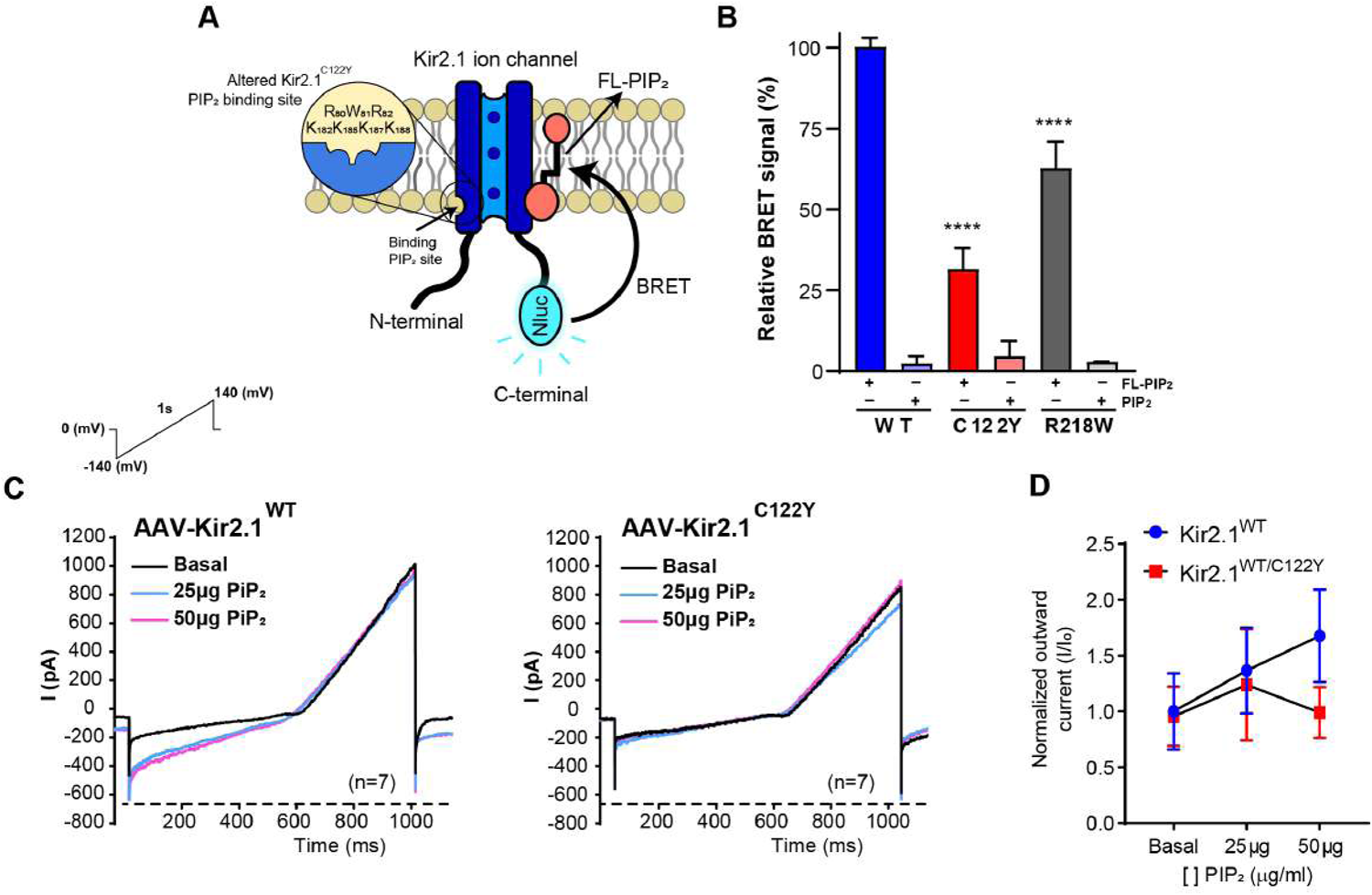
The C122Y mutation reduces Kir2.1-PIP_2_ binding capacity and interaction. **A**: Diagram of Kir2.1 monomer fused to the bioluminescent protein nanoluciferase (Nluc) (adapted from Cabanos et al.^25^). **B**: Specific BRET signal of binding Fl-PIP_2_ to Kir2.1 WT, C122Y and R218W and competition with non-fluorescent PIP2 version. Reduced binding was observed for C122Y and R218W (N=3 replicates per group; n=8-10 wells). **C**: Representative inside-out recording of I_K1_ in the absence (black current) and the presence of 25 (blue) and 50 (purple) μg/ml of PiP_2_. **D**: Normalized peak currents (I/I_0_) from −30 to +10 mV show that heterozygous condition abolishes the response to increasing PIP_2_ concentration. In contrast, in Kir2.1^WT^-transfected cells inward current increased progressively with PIP_2_. Both groups maintained an unaltered outward I_K1_. (n=7). Statistical analyses were conducted using two-tailed ANOVA. * p<0.05; ** p<0.01; **** p<0.0001

## Discussion

We report on the first human ATS1 mutation, C122Y, that breaks the Cys122-to-Cys154 disulfide bond in the extracellular domain of the tridimensional Kir2.1 structure. The disruption leads to defects in PIP_2_-dependent regulation, exerting a dominant negative effect with Kir2.1 tetramer channel dysfunction and life-threatening arrhythmias. Our AAV-mediated mouse model recapitulates *in-vivo* the ECG phenotype of the ATS1 patient carrying the C122Y mutation. ISO administration led to progressive further prolongation in the PR, QRS, and QT intervals. In addition, the mutation increases susceptibility to pacing-induced arrhythmogenic events of high severity (>1 second) in Kir2.1^C122Y^ animals relative to controls, including non-sustained ventricular tachycardias similar to those observed on the proband’s ECG. Isolated cardiomyocytes from Kir2.1^C122Y^ mice exhibited defects produced by decreased I_K1_ and I_Na_ compared to controls, including a a significantly depolarized RMP and reduced excitability. They also displayed prolonged APD, and in many cases EADs, bi-stability of the RMP and spontaneous calcium release events. The bistable resting membrane potential shown by some of the Kir2.1^C122Y^ cardiomyocytes may have been in part be due to the modification of the overall I_K1_ IV relation shape produced by the mutation. As demonstrated many years ago by Gadsby and Cranefield (1977) in Purkinje fibers, the existence of two possible stable resting potentials requires that the net steady-state IV relationship be “N-shaped,” with two zero-current intercepts in regions of positive slope conductance.^39^ A third unstable intercept occurs in a region of negative slope conductance. In the case of the Kir2.1^C122Y^ cardiomyocyte, the reduced Kir2.1 outward current at voltages between −60 and 0 mV, counterbalanced by the inward background conductance carried predominantly by sodium and calcium ions, generated an N-shaped current-voltage relation that crossed the voltage axis three times allowing two levels of resting membrane potential. Altogether, our results provide a potential mechanism for the spontaneous and induced arrhythmias observed in our ATS1 mouse model. While at baseline Ca^2+^ dynamics were similar to control after caffeine administration, ISO increased arrhythmia inducibility, suggesting abnormal Ca^2+^ dynamics transients.^24^.

Our *in-silico* homology modelling of the tridimensional Kir2.1 structure helps us understand the structural mechanisms underlying Kir2.1^C122Y^ dysfunction. Loss of the extracellular disulfide bond clearly alters the tridimensional structure and disrupts channel activity despite apparently normal Kir2.1^C122Y^ channel trafficking to the sarcolemma. However, despite channel reorganization, Kir2.1^C122Y^ still maintains a 78-84% similarity with Kir2.1^WT^ (TMscore: mutant heterotetramer: 0.7806 vs mutant homotetramer: 0.8391), which suggests a failure of Kir2.1^C122Y^ interaction with one or more key regulatory elements required for proper channel function. PIP_2_ signaling is a top candidate. PIP_2_ has emerged as a central subcellular mechanism for controlling ion channels and the excitability of nerves and cardiac muscle.^40^ PIP_2_ acts as a cofactor for proper Kir2.1 activity at the cell membrane. Kir2.1 channel-PIP_2_ interactions are crucial for channel activity and regulation, and defects in PIP_2_ binding constitute a major mechanism of Kir2.1 dysfunction underlying the loss-of-function in several ATS1.^9^

Our MD simulations with a single PIP_2_ molecule bound per monomer during 2000 ns MD replicas revealed that the mutation increased the probability of change in the Y_122_ Φ-dihedral angle leading to an altered hydrogen bond network in the extracellular loop. However, regardless of whether or not the dihedral angle varied, the distance between inner and outer helices remained unchanged. Nonetheless, the mutation triggered structural changes, particularly at the C-linker, which directly modified the PIP_2_ binding site comprising amino acids from two main structural regions of the channel. According to the Kir2.2 channel X-ray crystal structure (PDB code 3SPH), the 1′ PIP_2_ phosphate interacts with amino acids forming the sequence RWR (R_80_W_81_R_82_). This sequence is conserved (as RWR or KWR) among many different Kir channels and is located at the N-terminus of the outer helix.^35^ The RWR motif forms a binding site in which the 1′ phosphate caps the helix and is cradled by main-chain amide nitrogen atoms and the guanidinium groups of the two arginine residues. The tryptophan (W_80_) residue appears to anchor to the end of the outer helix at the membrane interface and also interact with one of the acyl chains. Similarly, 4’ and 5’ PiP_2_ phosphates interact and form hydrogen bonds with the helicoidal internal sliding helix (C-linker) at the end of the TM2 K_183_, K_186_, K_188_ and K_189_ residues in Kir2.2.^35^ Throughout the 2000 ns MD, hydrogen bonds between Kir2.1 and PIP_2_ decreased more rapidly for mutant channels compared to Kir2.1^WT^. Specifically, the PIP_2_ 1′ phosphate cap lost its interaction with the R_80_W_81_R_82_ triad, particularly R_82_, which appears strongly bound to WT monomers for longer simulation time (**Supplemental Table 2**).

PIP_2_ binding is known to induce a large conformational change in Kir channels leading to the formation of two new helices, an N-terminal extension of the ‘interfacial’ helix and a ‘tether’ helix at the C-linker.^35,36^ The flexible expansion of the C-linker contracts to a compact helical structure involving translation of the CTD ∼6Å towards the TMD, where it remains anchored and allows opening of the inner gate of the helix.^35,36, 45–47^ Importantly, separation between helices comes about as a result of slight splaying, but more significantly rotation of the inner helices, which moves hydrophobic amino acid side chains away from the ion pathway.^35^ Our MD simulation showed the C-linker disorganizing faster in Kir2.1^C122Y^ homo- and heterotetramer during the 2000 ns MD compared to WT channels, according to the loss of dihedral angles of P_186_ within those of the 3_10_-helix structure. These results highlight a rapid release of PIP_2_ molecules leading to channel closure, in accordance with the decreases in the Cα-Cα distance observed in the pore constriction residues Ile_176_, Met_180_ and A_306_, which also appeared barely dynamic over 2000 ns MD. In agreement, other studies have shown that P_186_ mutations lead to channel assembly, but with significantly reduced PIP_2-_ binding capacity.^41^ Taken together, these results suggest that C122Y induces a reorganization of the chains starting extracellularly and is transmitting along the channel to finally interrupts PIP_2_’s function. Nonetheless, the precise mechanism by which the C122Y mutation interferes with Kir2.1 binding to PIP_2_ molecules is beyond the scope of this study and remains to be fully elucidated.

Taking advantage of the BRET lipid binding assay, our results clearly show a significant decrease in the percentage of BRET signal in Kir2.1^C122Y^ channels, similar to the reduction in BRET signal for Kir2.1^R218W^ channels, with a well-known failure to interact with PIP ^10^ We next directly measured the functional effects of PIP_2_ on Kir2.1^WT^ and heterozygous Kir2.1^WT/C122Y^ channels in inside-out voltage-clamped membrane patches from transfected HEK293T cells. Altogether, the results showed that the C122Y mutation attenuated the maintenance of the I_k1_ current over time with increasing PIP2 concentration, which explained the lack of PIP_2_-dependent I_K1_ current. Thus, we validated the *in-silico* MD predictions and the demonstration by BRET that the Kir2.1^C122Y^ mutation breaks the disulfide bonds in the Kir2.1 extracellular domain, altering PIP2-dependent regulation to finally lead to channel dysfunction.

Interestingly, Macías et al. ^24^ have recently shown an SR microdomain of functional Kir2.1 channels contributing to intracellular Ca^2+^ homeostasis that could explain the phenotypic overlapping between ATS1 and catecholaminergic polymorphic ventricular tachycardia (CPVT) in some patients.^42,43^ Ca^2+^ fluxes across the SR membrane are bidirectional, and need a charge-compensating countercurrent ensuring that the SR membrane potential remains near 0 mV during the e-c coupling process.^44,45^ Importantly, our results demonstrate that intracellular Ca^2+^ homeostasis was similar in WT and C122Y under acute caffeine administration in intact isolated cardiomyocytes, suggesting that the SR Kir2.1 channel population is not regulated in a PIP_2-_dependent manner. However, a role for intracellular Ca^2+^ in arrhythmogenesis provoked by Kir2.1^C122Y^ was evidenced only after overloading the SR by ISO administration. The results suggest that while sarcolemmal Kir2.1^C122Y^ channels fail to conduct potassium through PIP_2_-dependent mechanisms, SR Kir2.1^C122Y^ channels remain functional independently of PIP_2_ activity. In support of such an idea, Katan et al. demonstrated that PiP_2_ is exclusively involved in sarcolemmal activities, including controlling Kir2.1 function.^46^

Kir2.1 channels are part of large multiprotein complexes comprising components of the cytoskeleton, regulatory kinases and phosphatases, trafficking proteins, extracellular matrix proteins, and even other ion channels.^47–49^ This probably explains in part the wide variety of clinical phenotypes found in different families with the same mutation and even within the same family.^3^ Kir2.1 forms channelosomes with Na_V_1.5, which indicates that the disease should no longer be considered in the simplistic terms of “monogenic” disorder.^2^ In fact, as our results show, it would not be correct to assume that the arrhythmic phenotype manifested by the patient is directly due to the mutation in question, but we must also consider potential modifications on the channeĺs interacting proteins. Therefore, the paradigm-shifting premise of this work is that we can no longer consider inherited arrhythmogenic diseases in terms of dysregulation of a single protein, because alteration of any member of a particular multiprotein complex has the potential to modify the function of associated proteins, resulting in a more complex disease. In this sense, the phenotypic manifestations in ATS1 are only understood by considering the wide range of proteins with which the altered ion channels interact.^49^ Our results show that Kir2.1^C122Y^ not only reduces I_K1_ but also I_Na_ in isolated mouse cardiomyocytes carrying the mutation. However, the C122Y mutation does not affect trafficking of ether Kir2.1 or Na_V_1.5 to the sarcolemma, suggesting new regulatory pathways for channelosome function. To further analyze molecular mechanisms involving Na_V_1.5 regulation, we studied channelosome homeostasis of both Kir2.1 and Na_V_1.5 proteins due to the differences in Gibbs free-energy values (WT: 4801.404 vs C122Y: −4131.754 for homo or −2274.207 for heterotetramer). Cardiomyocytes were treated with cycloheximide (CHX)^50^, a ribosomal RNA transcription inhibitor, for periods of 8, 16 and 24 hours at final concentrations of 100 µg/ml (**Supplemental Figure 10**). Interruption of protein synthesis resulted in a decrease of total Kir2.1 protein after 24 hours treatment compared to control **(Supplemental Figure 10A**). Immunostaining showed a significant co-localization with Rab5, protein involved in early endosomal formation (**Supplemental Figure 10B**), suggesting protein instability. Similarly, CHX decreased total Na_V_1.5 protein after 8 hours confirming the reduction in cell surface expression (**Supplemental Figure 10C**). However, regulation of Na_V_1.5 in these patients remains unclear. Reductions in Na_V_1.5 function/expression provide a slow-conduction substrate for cardiac arrhythmias. Van Bemmelen et al. demonstrate that Na_V_1.5 can be ubiquitinated in heart tissues and that the ubiquitin-protein ligase Nedd4-2 acts on Na_V_1.5 by decreasing the channel density at the cell surface^51^. Furthermore, in conditions like heart failure, elevated [Ca^2+^]_i_ increased Nedd4-2, interaction between Nedd4-2 and Nav1.5, and Na_V_1.5 ubiquitination with consequent degradation^52^, suggesting a crucial role of Nedd4-2 in Na_V_1.5 downregulation in heart disease. We looked for the expression in Nedd4-2 and our results showed a similar total expression of Nedd4-2 in both Kir2.1^WT^ and Kir2.1^C122Y^ cardiomyocytes, indicating that other regulatory pathways control the expression of Na_V_1.5 at the cell surface membrane as a consequence of the Kir2.1^C122Y^ mutation (**Supplemental Figure 10D**). Further studies are needed to elucidate the regulation of the Na_V_1.5 channel associated with the Kir2.1^C122Y^ mutation, but our data suggest a complex mechanism involved in channelosome function.

The potential clinical impact of this novel paradigm is groundbreaking: understanding the Kir2.1 modulation by its multiple interacting molecules will significantly improve our knowledge of channel function and of inherited and acquired arrhythmogenic cardiac diseases. It should also lay the groundwork for the generation of innovative, effective and safe approaches to prevent SCD in these and other devastating cardiac disease. Compared Together with previous data,^24^ all the results shown here support the hypothesis that the molecular mechanisms that increase the susceptibility to arrhythmias and SCD in ATS1 are different depending on the specific mutation, so that pharmacological treatment and clinical management should be different for each patient.

In conclusion, using AAV-mediated gene transfer we have generated a mouse model that recapitulated the electrocardiographic ATS1 phenotype of probands. ISO administration prolonged the PR, QRS and QT duration, and increased susceptibility to arrhythmogenic events of high severity. *In-silico* MD studies showed that the loss of the extracellular disulfide bond leads to channel closure by altering the Kir2.1-PIP_2_ hydrogen-bonding network. BRET and inside-out patch-clamping experiments confirmed the low ability of the Kir2.1^C122Y^ to properly bind PIP_2_ in sarcolemma. Altogether, this is the first demonstration that the break disulfide bond in the extracellular domain of the Kir2.1 channel results in defects in PIP_2_-dependent regulation, leading to channel dysfunction and life-threatening arrhythmias.

## FUNDING

Supported by National Institutes of Health R01 HL163943; La Caixa Banking Foundation under the project code LCF/PR/HR19/52160013; grant PI20 / 01220 of the public call “Proyectos de Investigación en Salud 2020” (PI-FIS-2020) funded by Instituto de Salud Carlos III (ISCIII); MCIU grant BFU2016-75144-R and PID2020-116935RB-I00, and co-funded by Fondo Europeo de Desarrollo Regional (FEDER); and Fundación La Marató de TV3 (736/C/2020): and CIBER de Enfermedades Cardiovasculares (CIBERCV), Madrid, Spain. We also receive support from the European Union’s Horizon 2020 Research and Innovation programme under grant agreement GA-965286; to JJ; Grant PID2021-126423OB-C22 (to MMM) funded by MCIN/AEI/ 10.13039/501100011033 and by “ERDF A way of making Europe”; Grant PID2019-104366RB-C22 (to MGR), funded by MCIN/AEI/10.13039/501100011033; Funding provided by the Dynamic Microscopy and Imaging Unit - ICTS-ReDib Grant ICTS-2018-04-CNIC-16 funded by MCIN/AEI /10.13039/501100011033 and ERDF; project EQC2018-005070-P funded by MCIN/AEI /10.13039/501100011033 and FEDER. CNIC is supported by the Instituto de Salud Carlos III (ISCIII), the Ministerio de Ciencia e Innovación (MCIN) and the Pro CNIC Foundation, and is a Severo Ochoa Center of Excellence (grant CEX2020-001041-S funded by MICIN/AEI/10.13039/501100011033). AIM-M holds a FPU contract (FPU20/01569) from Ministerio de Universidades. LKG holds a FPI contract (PRE2018-083530), Ministerio de Economía y Competitividad de España co-funded by Fondo Social Europeo ‘El Fondo Social Europeo invierte en tu futuro’, attached to Project SEV-2015-0505-18-2. IMC holds a PFIS contract (FI21/00243) funded by Instituto de Salud Carlos III and Fondo Social Europeo Plus (FSE+), ‘co-funded by the European Union’. MLVP held contract PEJD-2019-PRE/BMD-15982 funded by Consejería de Educación e Investigación de la Comunidad de Madrid y Fondo Social Europeo ‘El FSE invierte en tu futuro’.

## DISCLOSURES

None

## ACKNOWLEDGEMENTS

We thank the CNIC Viral Vectors Unit for producing the AAV9. The confocal experiments were carried out in the CNIC Microscopy and Dynamic Imaging Unit. We thank the CNIC Bioinformatics Unit for generating the *in-silico* homology modelling simulations, F-function analysis and helping in their discussion. We also thank the Centro de Supercomputación de Galicia (CESGA) for use of the Finis Terrae III supercomputer to perform molecular dynamics studies.

## AUTHOR CONTRIBUTION

F.M.C. and J.J. co-designed the experiments; F.M.C. performed most of the experiments; A.M. and A.I.M.M are author for cellular electrophysiology; A.D., E.Z., and J.J.J. provided clinical data; L.K.G. supported optical mapping experiments: L.K.G. and F.M. were in charge of *in-silico* homology modelling and molecular docking studies; A.D.A, M.M.M. and M.G.R. performed molecular dynamics simulation and analysis and provided funding; M.L.V., P.S.P., J.R.R., S.A.S., I.M.C., F.B.J., A.B.B. and J.A.B. provided technical support, discussions and revisions; F.M.C. and J.J. co-wrote the manuscript and conceived the study; J.J. provided supervision, funding and revisions; All authors discussed the results and commented on and approved the manuscript.

## CORRESPONDING AUTHORS

Correspondence to José Jalife or Francisco M. Cruz.

## SUPPLEMENTAL INFORMATION

Extended Materials and Methods

Supplementary Figures 1-10

Supplementary Tables 1-4

## Notes

### Competing Interest Statement

The authors have declared no competing interest.

